# Age Dependent Integration of Cortical Progenitors Transplanted at the Adult CSF-Brain Interface

**DOI:** 10.1101/2024.08.28.609860

**Authors:** Nikorn Pothayee, Gretchen Greene, Jahandar Jahanipour, Hyesoo Jie, Jung-Hwa Tao-Cheng, Emily Petrus, Dragan Maric, Alan P. Koretsky

## Abstract

There has been renewed interest in neural transplantation of cells and tissues for brain repair. Recent studies have demonstrated the ability of transplanted neural precursor cells and in vitro grown organoids to mature and locally integrate into host brain neural circuitry. Much effort has focused on how the transplant behaves and functions after the procedure, but the extent to which the host brain can properly innervate the transplant, particularly in the context of aging, is largely unexplored. Here we report that transplantation of rat embryonic cortical precursor cells into the cerebrospinal fluid-subventricular zone (CSF-SVZ) of adult rat brains generates a brain-like tissue (BLT) at an ectopic site. This model allows for the assessment of long-range connectivity and cellular interactions between the transplant and the host brain as a function of host age. The transplanted precursor cells initially proliferate, then differentiate, and develop into mature BLTs, which receive supportive cellular components from the host including blood vessels, microglia, astrocytes, and oligodendrocytes. There was integration of the BLT into the host brain which occurred at all ages studied, suggesting that host age does not affect the maturation and integration of the transplant-derived BLT. Long-range axonal projections from the BLT into the host brain were robust throughout the different aged recipients. However, long-distance innervation originating from the host brain into the BLT significantly declined with age. This work demonstrates the feasibility and utility of integrating new neural tissue structures at ectopic sites into adult brain circuits to study host-transplant interactions.

## Introduction

Neurons of the central nervous system (CNS) generally lack the ability to repair and regenerate following injury or disease (Harel & Strittmatter, 2006; Varadarajan et al., 2022). Cell and tissue transplantation aimed at forming a defined brain tissue presents a promising strategy for repairing and restoring brain function in individuals affected by neurodegenerative diseases and brain injuries (Barker et al., 2018; Grade & Götz, 2017). Early studies on fetal tissue transplantation demonstrated the ability of embryonic neuronal tissue to survive and form functional connections when implanted into young rodent brains (Gash et al., 1985; Girman & Golovina, 1990; Lu & Norman, 1993). Transplantation of more defined neuronal cell grafts and neuronal progenitor cells (NPCs) showed the implanted cells differentiate into mature neurons capable of extending local and long-range projections in the brain and spinal cord (Denham et al., 2012; Falkner et al., 2016; Gaillard et al., 2007; Lu et al., 2012; Michelsen et al., 2015; Pothayee et al., 2018). In addition to efferent projections from the graft or transplant derived tissue, some studies have shown functional integration of the new tissue through the establishment of local host-transplant synapses (Espuny-Camacho et al., 2018; Falkner et al., 2016; Kumamaru et al., 2019; Michelsen et al., 2015; Palma-Tortosa et al., 2020; Pothayee et al., 2018). Recently, advances in the development of 3D neural organoids have demonstrated the ability to generate autologous complex brain tissues and numerous studies have successfully transplanted human neural organoids into rodent brains (Jgamadze et al., 2023; Kelley et al., 2024; Kitahara et al., 2020; Mansour et al., 2018; Revah et al., 2022; Velasco et al., 2019; Wang et al., 2024). Both organoid and cell transplantations offer promising avenues for repairing neural circuits and studying the mechanisms of postnatal circuit formation.

The majority of transplantation studies have demonstrated the maturation and integration of neural transplants – either organoids or cell-derived implanted into newborn or critical period, young host brains. It remains unclear if high level integration and graft-host cellular interaction can occur in aged animals. For example, it is unclear whether a new, long-range axonal projections that innervate the transplant from distant host brain sites can form in aged animals. We reported previously that the CSF environment enabled early neural precursor cells to grow into brain-like tissues (BLTs) without signs of teratomas or cancerous tissue (Pothayee et al., 2018). Extensive integration of host cells and axons projecting from the BLT into the host were observed. The ectopic site of tissue growth makes this an attractive model to study graft-host integration and new circuit formation. However, a limitation of the previous study included the nonspecific location of the BLT’s site of implantation. In this study, the BLT was grown near the SVZ, close to the rostral migratory stream (RMS). The ability of transplanted cells to mature, send projections into the host brain, and recruit specific long-distance innervation from the host was determined.

Aging negatively impacts new cell and circuit formation (Babcock et al., 2021; Couillard-Despres et al., 2011). Therefore, we hypothesized that increasing age of the host may impair survival and integration of the BLT with host circuitry. Here we tested the limits of aging’s impact on these factors by studying the cellular and neuronal functional integration of the BLT with the host at varying ages. Using this approach, we found that the host age up to one year does not limit or alter growth potential and phenotype of the BLT or the host contributions to the BLT such as blood vessels and microglia. However, increasing recipient age decreased the prevalence of long-range connections from the host brain projecting into the BLT. Despite this decrease, new projections into the BLT from the host could still be detected in one year old host animals.

## Results

### 1. Formation and integration of neural precursor cells transplanted at the CSF-SVZ interface

Freshly sorted lineage negative cortical neural precursor cells were isolated from primary GFP-expressing E13.5 rat embryonic dorsal telencephalic tissues as previously described (Maric et al., 2007; Pothayee et al., 2018) (Figure 1A, Extended Figure 1). These cells were injected in the SVZ adjacent to the RMS in adult rat brains after isolation. After eight weeks post-implantation (PI), the transplanted cells developed into a stable brain-like tissue (BLT) which was detected with T_2_-weighed MRI (Figure 1B). These transplanted cells initially exhibited high proliferating (PCNA-positive) activity within one week PI, but proliferation subsided by eight weeks after implantation (Extended Figure 2). Immunostaining with NeuN showed an abundance of mature neurons in the transplant-derived BLT. Approximately 80% of the NeuN positive cells in the BLT were GFP-positive, indicating that they were derived from the injected neural precursor cells. The remaining 20% of NeuN positive cells in the BLT were GFP-negative, indicating that they originated from the host (Extended Figures 3, Figure 2C), in agreement with our previous study (Pothayee et al., 2018).

**Figure 1.**
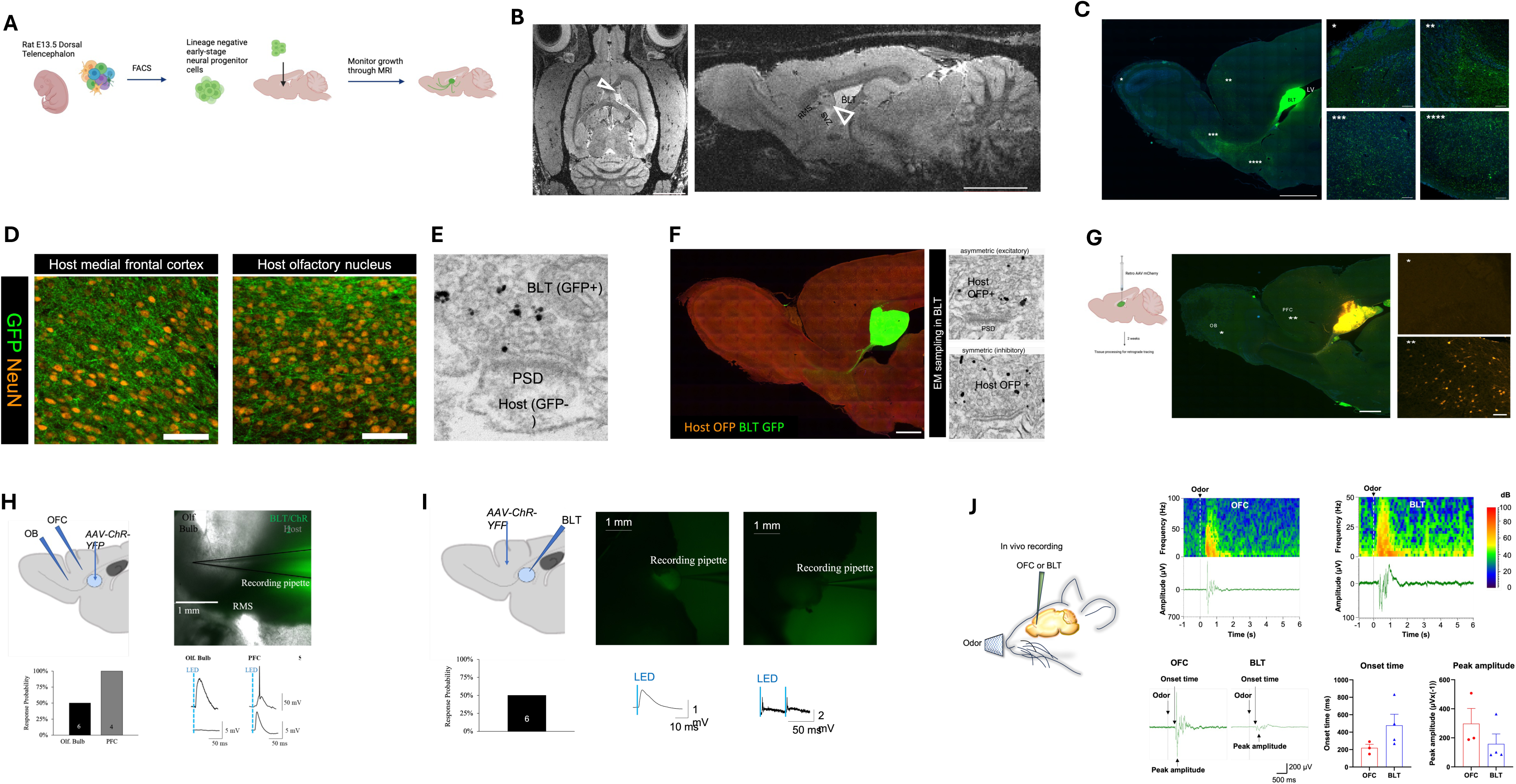
Formation and integration of brain-like tissue at of the CSF-SVZ-RMS interface. **A**. Workflow of cell implantation. Embryonic rat cortical neural precursor cells were isolated from dissociates of E13.5 dorsal telencephalic tissue and purified FACS prior to implantation into the subventricular zone (SVZ) next to rostral migratory stream (RMS). **B.** Axial and sagittal MRI of rat brain eight weeks post-implantation (PI), at which point the precursor-generated brain-like tissue (BLT) transplants were clearly visible (open arrowheads). Scale bar = 2 mm. **C.** Sagittal brain section showing GFP-postive BLT (green) near the SVZ/RMS injection site bordering the CSF-filled lateral ventricle (LV). The BLT projects into various regions of the forebrain** and olfactory system (*, **, ***) Scale bar = 2 mm (larger FOV) and 100 um (smaller FOV) **D.** High-magnification fluorescent images of neural processes from the GFP-positive BLT innervating the host neurons (NeuN-positive cells) in the host front cortex and host olfactory nucleus. Scale bar = 100 um. **E.** Electron micrograph of the presynaptic terminal from BLT (GFP-postive) that forms a synapse with the host (GFP-negative) neuron. A prominent postsynaptic dendity (PSD) indicates the excitatory nature of this synapse. **F.** A grafted BLT (green GFP-positive) integrating into a host brain (orange OFP-positive). Electron micrographs showing asymmetric (excitatory) and symmetric (inhibitory) synapses from different OFP-positive host axons innervating onto the OFP-negative BLT denritic spine and shaft, respectively. **G.** Retrograde tracing with AAV from the BLT shows projections into host prefrontal cortex but not olfactory bulb. Scale bar = 1 mm. **H.** Slice electrophysiology detects functional, monosynaptic connections from the ChR2-expressing BLT and the host medial prefrontal cortex and olfactory bulb. **I.** Slice electrophysiology shows synaptic connection from the ChR2-expressing host neurons in medial prefrontal cortex to the BLT. Note that some connections were strong and yielded action potentials (H, PFC), while others were weaker and only produced depolarizations (H, olfactory bulb). **K.** In vivo recording of rat with BLT was completed using 10% amyl acetate diluted in mineral oil as stimulus.. The left panel shows a schematic diagram of in vivo extracellular recordings using a 32-channel silicon probe. The top right panel shows LFP power spectrogram during odor stimulation in the mPFC (left) and BLT (right). The bottom shows example raw traces of LFP signals. Black arrows indicate onset time and peak amplitude. Onset time was compared between the mPFC group and the BLT group (mPFC: N = 3, BLT: N = 4). t-test, p > 0.05. Mean ± SEM. Peak amplitude was compared between the two groups (mPFC: N = 3, BLT: N = 4). t-test, p > 0.05. Mean ± SEM.

**Figure 2.**
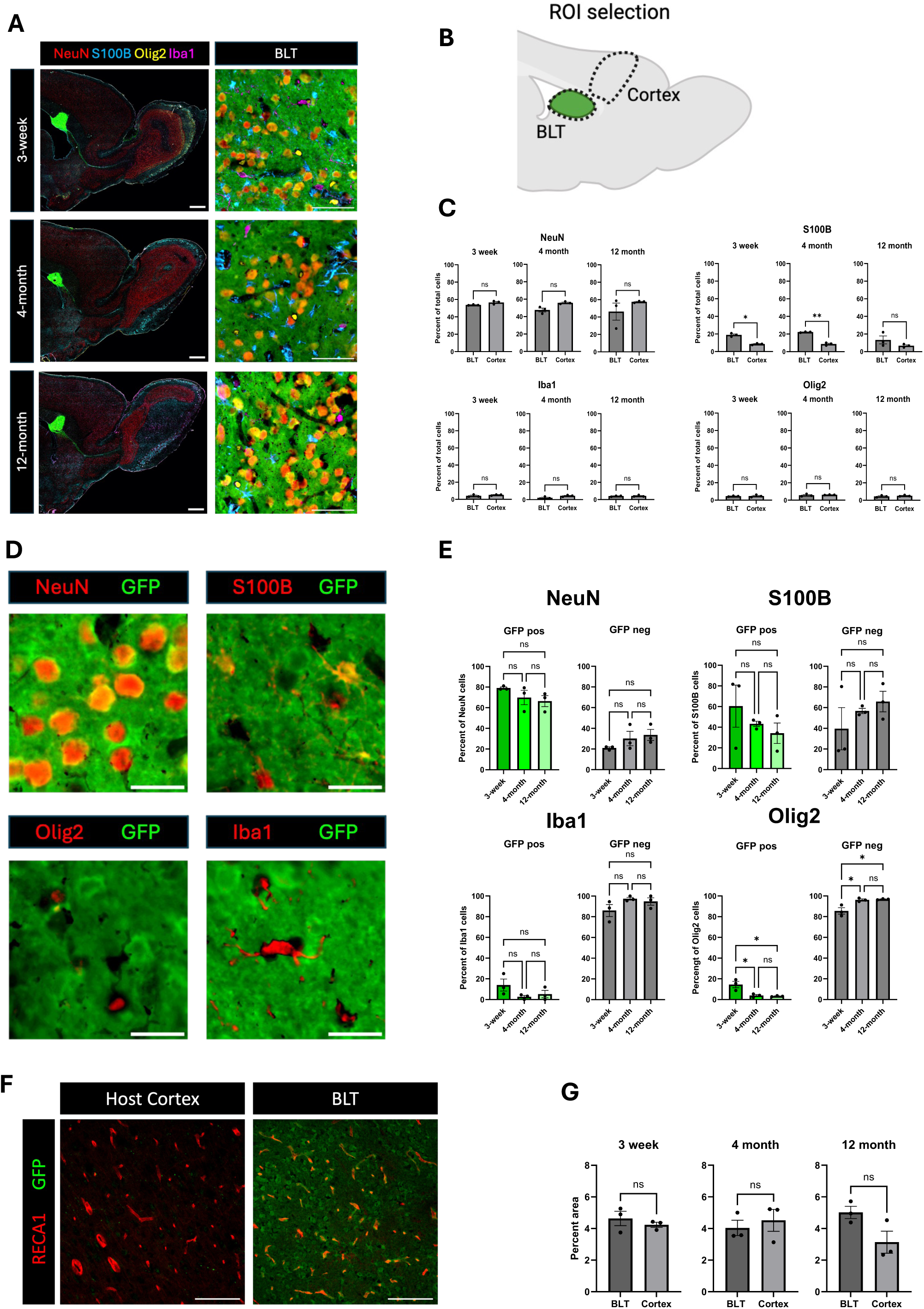
Transplanted neural progenitor cells form brain-like tissue closely resembling host cortex in three-week, four-month, and twelve-month old recipient rats. **A.** Fluorescent images of sagittal brain sections collected eight weeks after cell implantation into three-week, four-month, and twelve-month-old rats. At time of euthanasia, the animal were approximately three-month, five-month, and fourteen-month-old, respectively. NeuN-positive cells represent neurons. S100B-positive cells represent astrocytes, Iba1-positive cells represent microglia, and Olig2-positive cells represent oligodendrocytes. Neurons, astrocytes, microglia, and oligodendrocytes were found in the transplant across the age groups. **B**. Regions of interest (ROIs) for comparison of the cellular composition between the transplant and the host cortex (Created with BioRender.com). **C.** Comparison of cellular composition of the BLT and host cortex based on the percentage of each cell type relative to the total cells in the defined ROI (N = 3). Normality was evaluated and then an unpaired T-test was used two compare the two conditions. NS means no statistical difference was observed. *** show p value = 0.001. **D.** Fluorescent images from immunostaining characterizing origin of each cell type. GFP positive cells signify transplant origin, whereas GFP negative cells are derived from the host. **E.** Percent of each cell type – neurons, astrocytes, microglia, and oligodendrocytes – derived from implanted cells (GFP-positive) vs host (GFP-negative) compared between the three age groups. A one-way ANOVA followed by Tukey’s multiple comparisons test was used. **F.** Immunostaining with RECA-1 for identification of blood vessels. **G.** Vasculature density in the transplant and host cortex across the host ages (N= 3). Normality was evaluated and then an unpaired T-test was used to compare the two conditions.

Next, we determined whether the BLT integration with the host brain included sending long-distance connections into the host. Immunostaining with anti-GFP antibodies showed that the transplant extended projections from its implantation site into the host brain (Figure 1C). These projections were predominantly located along the RMS into the olfactory bulb. In addition, GFP-positive projections from the BLT were observed in the hosts’ frontal cortex, striatum, and thalamus (Figure 1C insets, 1D). Immuno-gold staining and electron microscopy (EM) showed that indeed these GFP-positive projections from the BLT formed synapses with GFP-negative host neurons located outside the BLT (Figure 1E). Innervation from the host brain into the transplant was also observed by implanting GFP-positive neural precursor cells into host rodent brain that ubiquitously expressed orange fluorescent protein (OFP) (Figure 1F). EM immunogold showed the presence of both asymmetric (excitatory) and symmetric (inhibitory) OFP-positive presynaptic terminals located adjacent to OFP-negative neurons (implanted cell-derived) (Figure 1F). These results indicate that the BLT forms multiple long-range connections with the host. Retrograde tracing from the BLT showed host innervation originating from the medial frontal cortex and thalamus. No retrograde AAV labeling was observed in the olfactory bulb of the host even though the BLT sent projections to the olfactory bulb (Figure 1G).

Slice electrophysiology was used to examine whether the connections between the neural transplant and the host were functional. An adeno-associated virus encoding for ChR2 expressing a yellow, fluorescent marker (AAV-ChR2-eYFP) was injected into the BLT approximately eight weeks after implantation. Six weeks after AAV transfection host neurons in the olfactory bulb area and medial frontal cortex were patched for electrophysiological recordings. Light was used to activate ChR2-expressing synapses from the transplant and monosynaptic connections were detected in both areas. Between 50-100% of host neurons in areas innervated by the BLT responded to light stimulation (Figure 1H). To determine if the projections from the frontal cortex were active in the BLT, ChR2-AAV was injected into the host frontal cortex and neurons in the transplant were patched for recordings. We found that 50% of the cells in the BLT responded to light stimulation of host-innervated regions (Figure 1 I). These results demonstrate that bidirectional synaptic activity between the host and the transplant were functional. Finally, we assessed whether the neural transplant responded to external stimuli in vivo. We used odor stimulation because anterograde and retrograde tracing detected BLT integration into the olfactory system and frontal cortical regions that are involved in odor processing. A pulse of 10% amyl acetate scented air elicited robust local field potential (LFP) response, approximately 200 μV, in the BLT in 4 host rats (Figure 1K, middle panels). The LFP response from recordings at the medial frontal cortex of the host rat brain responded with a shorter latency by approximately 300 ms, indicating that the route of input from the olfactory bulb may be via frontal cortex.

### 2. Impact of host age on neural transplant development

The interaction between host and the BLT are complex and involve multiple glial cell types (Talaverón et al., 2014). Microglia, vessels, and most oligodendrocytes migrated into the BLT from the host, while most neurons and astrocytes arose from the BLT. To evaluate the impact of host age on the BLT’s maturation, we implanted early-stage NPCs into the SVZ region of rats that were three-weeks, four-months, or 12-months of age. Eight weeks following implantation, host rats were euthanized, and the cellular composition of the BLT was evaluated using multiplexed immunohistochemistry with automated cell counting (Maric et al., 2021). The implanted cells developed into a similar ratio of neural cell phenotypes regardless of the age of the recipient. Using NeuN to label mature neurons, Olig2 for oligodendrocytes, Iba1 for microglia, and S100B for astrocytes, we compared the overall cellular composition of the BLT to normal frontal cortical tissue located distal from the cell implantation site (Figure 2A and 2B). The percentage of neurons, oligodendrocytes, and microglia of the total cells enumerated within both regions was not significantly different irrespective of transplant recipient host age (Figure 2C). In contrast, the percent of total astrocytes was significantly higher in the BLT compared to the host frontal cortex in the 3-week-old (19% in BLT vs 8.6% in cortex, p value = 0.0195) and 4-month-old group (22.17 % in BLT and 8.7% in cortex, p value =0.0075), but not significantly different in the 12-month-old group (p value = 0.2319). These findings imply that there was increased astrocyte number in the BLT relative to the cortex in younger hosts. The distribution of BLT-derived and host derived cells was measured by co-staining the GFP-positive BLT tissue with known cell markers (Figure 2D). Within the BLT, the implanted precursor cells primarily matured into neurons (GFP-positive and NeuN-positive) and secondarily into astrocytes (GFP-positive and S100B-positive) across all three age groups (Figure 2D, E). Most oligodendrocytes and microglia were GFP-negative (Figure 2D, 2E) and were thus supplied by the host. This agrees with previously reported data in younger rodents and follows the known differentiation potential of the cells isolated from the early (E13.5) dorsal telencephalon, which at that gestational stage do not generate oligodendrocyte precursors (Su et al., 2023). If any microglia infiltrated the diencephalon at E13.5, they were presumably depleted by lineage negative selection using FACS.

In all hosts, the BLT was well vascularized as demonstrated by immunostaining for Rat Endothelial Cell Antigen (RECA)-1 (Figure 2F, Extended Figure 4). There was no significant difference between the density of blood vessels in the BLT in comparison to the host cortex in any of the age groups (Figure 2G, three-week: p value= 0.4542, four-month: p-value= 0.6035, 12-month: p-value= 0.0779). The vessels were host derived, and angiogenesis occurred in the BLT despite the advanced age of the host. Further evaluation of markers such as myelination (MBP), inhibitory neurons (GAD67), and the formation of a host ependymal layer at the interface of BLT-to-CSF were comparable across all age groups as well (Extended Figure 5-7).

### 3. Long-range connectivity between BLT and host as function of age

Long-range connectivity between neural transplants of both cells and organoids is of great interest in order to establish functional circuits that are integrated into the host brain. Here, we sought to determine whether recipient age impacts the ability of the transplant and host to establish such connections. BLT neurons project into the host circuitry extending into the olfactory bulb, medial prefrontal cortex, and the striatum (Figure 3A and 3B). The percentage of GFP fluorescent puncta was highest in the olfactory bulb and striatum with densities (i.e., total pixel area) of approximately 2% in these regions and less in the prefrontal cortex with a density of about 1%. Fluorescent puncta were also noted in the thalamus and piriform cortex (Figure 3B and Figure 3C, Extended Figure 8). There was no significant difference of transplant innervation from the BLT into the olfactory bulb, striatum, and PFC across the three host age groups (Figure 3C; olfactory bulb: p value = 0.4428, striatum p value = 0.3116, PFC: p value =0.8179). There was a significant difference seen with BLT innervation to the dorsal thalamus between the three-week and four-month-old groups (p value = 0.0476) with an increase in innervation in the four-month-old host. There was no difference seen in thalamic innervation between the three-week and 12-month (p value = 0.3138) or the four-month and 12-month hosts (p-value =0.3545). Therefore, the increase in innervation seen in the dorsal thalamus region between the three-week and four-month host might be due to the overall small percent area occupied by the fluorescent puncta and not reflect a true biological change in BLT innervation between the two host ages. Remarkably, the transplanted neurons are effective at sending projections into all host age groups.

**Figure 3.**
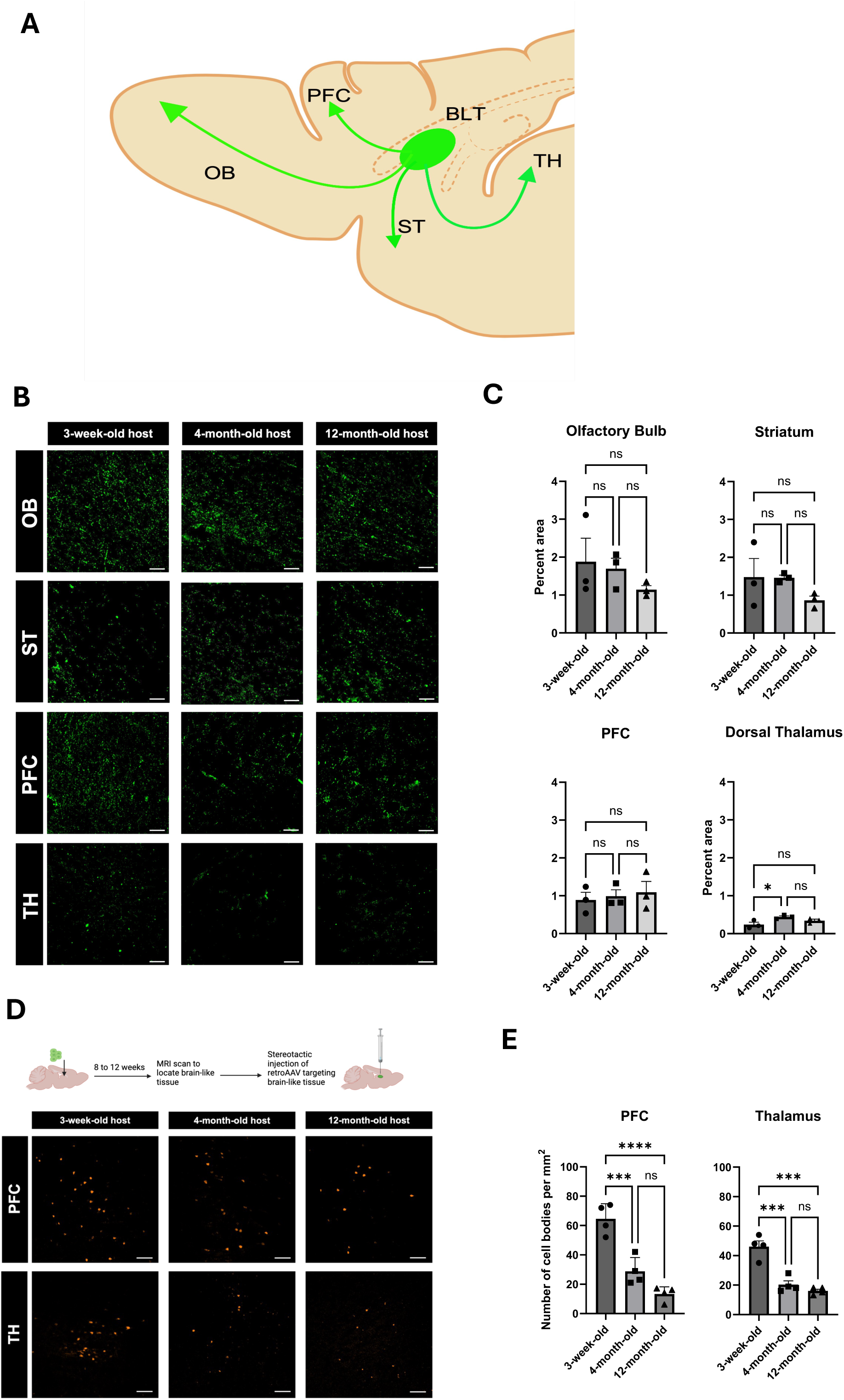
Impact of host age on bidirectional graft-host innervation. **A.** General diagram of BLT projections into different host brain regions including the olfactory blub (OB), striatum (ST), medial prefrontal cortex (PFC), and thalamus (TH) (Created with BioRender.com). **B** Afferent axonal projections from the BLT into the host olfactory blub, striatum, medial prefrontal cortex, and striatum in all recipient age groups. **C.** Quantification of afferent axonal projections based on the percent area of the fluorescent puncta out of total area compared across the three age groups (N = 3). Analysis was completed with a one-way ANOVA and Tukey’s multiple comparisons test. **D.** Retrograde AAV tracing from the BLT. Cell bodies of host neurons labeled by retroAAV-mCherry in prefrontal cortex and thalamus. **E.** Number of cells retrogradely labeled in each ROI compared between the three recipient age groups (N = 3). A one-way ANOVA with a Tukey’s multiple comparisons test was used for the comparison.

Next, we assessed the long-distance input originating from the host brain and projecting into the transplant using retrograde viral tracing. After injecting mcherry-expressing retrograde AAV into the BLT, we found labeled neuronal cell bodies located in the medial prefrontal cortex and the thalamic areas (Figure 3D) indicating that the new tissue received input from these host regions. However, there was a progressive decline in the number of neurons that projected into the transplant as the host age increased (Figure 3E, PFC: three-week vs four-month p-value = 0.0006, three-week vs 12-month p-value= <0.0001, thalamus: three-week vs four-month p-value = 0.0003, three-week vs 12-month p-value= 0.0001). No retrograde labeling was observed in the host olfactory bulb, striatum, and piriform cortex at any host age. There was a reduction of BLT innervation by the host neurons which was more robust with increasing age. Comparing the three-week-old host to four-month-old host, there was approximately a 55% decline of BLT innervation by the host neurons, and with a 12-month-old host, an almost 70% reduction in these numbers was identified. Overall, the host’s age did not limit neuronal innervation from the BLT to the host brain but did have a negative impact on the host’s ability to innervate the BLT.

## Discussion

There is renewed interest in tissue and cell implants into the central nervous system due to the development of brain organoids and the ability to differentiate induced pluripotent cells to specific neuronal cells from patient derived cells for transplantation (Jgamadze et al., 2023; Quezada et al., 2023; Velasco et al., 2019). Most studies to date have focused on the ability of the implant to survive, mature, and integrate into local circuits following transplantation (Dulin et al., 2018; Espuny-Camacho et al., 2013; Falkner et al., 2016; Gaillard et al., 2007; Gronning Hansen et al., 2020). Few studies have evaluated if and how the host brain adapts to accept the transplant. In previous work we showed that cortical precursor cells transplanted at the CSF-tissue interface can develop into a brain-like tissue with extensive interactions with the existing host brain tissue (Pothayee et al., 2018).

In the present study, we demonstrate the specific source of the host’s long-range connections with the BLT. By growing the BLT at an ectopic site at the CSF-tissue interface, this approach allows for the ability to quantitate host-BLT interactions and reduce local neural interactions between the host and graft. Based on this principle, our aim was to determine how the age of the host brain environment affects maturation, integration, and further interactions between the cortical precursor cells and the host brain circuitry.

### Cellular Composition of the BLT and Host Contribution to BLT are not affected by host age

Extensive studies have been done on transplanting cells, fetal tissue, and organoids into different parts of the brain and spinal cord in either intact or damaged conditions (Espuny-Camacho et al., 2013; Falkner et al., 2016; Gaillard et al., 2007; Girman & Golovina, 1990; Kitahara et al., 2020; Lu et al., 2012). In this study, we implanted neural precursor cells at the CSF-SVZ brain interface near the RMS. This location is known for its ability to generate new neurons throughout life, thus offering a permissive environment for neural precursor cell survival, proliferation, and differentiation (Fuchs et al., 2004). This differs from our previous report in which these cells were injected directly into the CSF in the lateral ventricle and formed a BLT inside the ventricle cavity. Though they productively incorporated with the host brain tissue in this earlier study, there was less control over where that occurred and multiple tissues were formed in the CSF (Pothayee et al., 2018). Nonetheless, these earlier studies showed extensive projections from the transplant to the host and some host innervation of the BLT. The growth of cells injected close to the SVZ as done in the present study followed the same pattern of proliferation and differentiation previously observed with injection to the CSF. The proliferation and differentiation of implanted cells progressed for about eight weeks post-injection, at which point the transplant-derived BLT no longer increased in size. These findings suggest that despite using a different injection site, the implanted neural precursor cells maintained the same proliferation and differentiation potential as previously observed.

The present study evaluated how recipient age impacts the ability of the implanted precursor cells to mature and form a tissue integrated with the host brain. No study to date has evaluated how aging might impact the ability of transplanted cells, organoids, or tissues to survive and mature. This question is particularly relevant given the goal of translating tissue transplantation to patients. Neurodegeneration, stroke, and other brain injuries primarily occur in adults, and the number of patients impacted increases with age (Feigin et al., 2020). Therefore, it is critical to understand how the adult and aged host environment might impact neural precursor cell survival and maturation. In all age groups, we observed that implanted NPCs can form a complex brain-like tissue closely resembling the cellular composition of the host cortex. The abundance of neurons and glia in the BLT was similar to the proportions seen in the normal cortex. While we observed increased astrocyte numbers in the BLT compared to the young host cortex, the values in older animals were not significantly different. This increase could be because the young host brain environment promotes astrocyte differentiation (Akdemir et al., 2020). Indeed, astrocyte ratio in the cortex of the young host studied is relatively lower compared with the older hosts.

Regardless of the age of the recipient, the host was able to vascularize the graft and supply microglia and oligodendrocytes. Additionally, an ependymal lining formed along the interface of the graft and CSF in all age groups, further demonstrating the seamless incorporation of the transplant derived tissue with the host. Overall, the ability of the transplanted NPCs to form a tissue structure closely resembling the host brain demonstrates the feasibility of cell transplantation even in aged host environments.

### The BLT sends and receives functional connections

Functional and reciprocal connections were observed between the BLT and the host brain. Axons extended from the BLT into the host forebrain and olfactory system as well as into the thalamus and adjacent striatum. While the mechanisms regulating the path by which the transplant derived BLT extended projections into the host brain are not yet characterized, it is possible that the existing host brain tissue provided guidance cues for axonal growth from the newly formed BLT into the existing brain neuronal circuitry. For example, projections into the olfactory bulb from the BLT tended to follow the nearby rostral migratory stream. However, it is not clear why the BLT sent projections to thalamus or frontal cortex especially in the older animals. The proximity of the striatum to the BLT could have facilitated projections to this region.

In previous work, there was evidence of the host extending axons into the BLT, but the origin of these host connections was not evaluated. Using retrograde AAV tracing, we showed that the host brain innervated the BLT mostly from the medial prefrontal cortex in the forebrain region and to a lesser extent the thalamus. Despite extensive innervation from the BLT into the host olfactory system, no reciprocal connections originating from the olfactory bulb or olfactory cortex were observed. It is believed that all long-distance connections have formed in the rodent by two to three weeks after birth (Price et al., 2006). Therefore, it will be important to determine how these projections from the host into the BLT form. It could be that these connections are due to axons shifting from their normal projections into the BLT, or that the BLT induces bifurcation of existing projections, least likely is the possibility that the BLT induces new axons from the host to innervate this newly-formed tissue. During rodent development, for example when the thalamo-cortical connections are made in somatosensory cortex, the thalamic connections extend to the cortex in coordination with the reciprocal cortical connections (Molnár & Blakemore, 1995). Therefore, it seems likely that the BLT extended axons into brain areas in coordination with projections that were sent into the BLT from the host. The difference in maturation and plasticity between different brain regions could determine where the BLT projects and which regions project back into the BLT. For example, the frontal cortex is known to have a later critical period then olfactory bulb. However, the observation that the host projected into the BLT even in one year old animals indicated that the BLT was able to recruit projections even from older host animals.

We detected synaptic connections between the BLT and the host. These were determined to be mature and could be involved for processing olfactory stimuli. The small response of the BLT to pure air or mock odors raises the possibility that the BLT was sensitive to the hosts’ arousal state. Both odor and arousal likely arrive via connections from the host to prefrontal cortex and may include neuromodulatory axons such as those containing epinephrine or acetylcholine. Future studies can be designed to evaluate if the BLT impacts tasks related to odor detection or discrimination.

The observation of long-distance connections from the BLT into the host is consistent with previous transplant studies which have shown that these projections typically form in young host brains. Here we show for the first time that age does not impair this ability to form connections. Remarkably, increasing host age up to one year of age did not affect the ability of the BLT to send projections to the host. These data suggest that the BLT maintains the ability to develop and extend neural outgrowths despite an aged host environment.

### Importance of Host Age on Long-distance Innervation of Host into Transplants

There are many examples now of neural transplants including cells, fetal tissue, and human cerebral organoids extending long distance neural processes into the host brain. Transplanted cells have been shown to functionally integrate into host circuitry (Dulin et al., 2018; Espuny-Camacho et al., 2018; Falkner et al., 2016; Kumamaru et al., 2019; Michelsen et al., 2015; Palma-Tortosa et al., 2020). There has been less work determining whether the host innervates the transplant over long distances. Some exceptions have shown longer distance connectivity from the host to the transplant, for example in the spinal cord (Lu et al., 2012). Recently, numerous studies have successfully demonstrated the ability of human neural organoids to survive and mature after implantation into host rodent brains between neonatal to three-month-old hosts (Jgamadze et al., 2023; Mansour et al., 2018; Revah et al., 2022). However, most studies have not determined whether connectivity arises from local connections with host tissue on the border of the transplant versus from a long-distance projection. The impact of host age on connectivity between the human-neural organoids or cells and the rodent host brains remains unclear. Indeed, to the best of our knowledge, 6-month-old mice are the oldest host animal used for cell implantation where subsequent cell maturation has been shown (Gaillard et al., 2007).

In this study, we sought to address this issue by placing the transplant ectopically to the host brain tissue, thus minimizing innervation from existing local connections. Using this approach, we have shown that increasing host age correlated with reduced long-distance projections from the host frontal cortex and thalamus into the transplant-derived BLT, with the most pronounced reductions in 12-month-old host rats. However, the presence of minimal but detectible projections implies that making these connections is feasible. It will be interesting to test whether this decline in connection could be rescued.

### Limitation and technical considerations of the study

Attempts were made to control the location of the transplanted cells, though it is still relatively difficult to ensure that the implanted cells do not spread throughout the CSF. In some cases, when this happened, we did not observe detectable tissue located near the injection site. Possible ways to ameliorate this issue would be through the implantation of preformed tissues or cells embedded in a 3D matrix. In addition, it is unknown if human neural organoids could be successfully implanted and grown in the SVZ-CSF rat host brain with similar results to matched homotopic NPC-host species. While we show that aging negatively impacts the ability of the host brain to form long-distance connections with the NPC-derived BLT, it’s not clear whether advanced aging (e.g., 18-24 months old rats or mice) would still be permissive for generation of new long-distance connections from the BLT into the host brain and vice versa. In addition, this present study utilized transplanted cells from an early developmental stage which likely facilitates a robust host response and a greater incorporation of the transplant to the host circuitry. Additionally, rat cortical precursor cells were transplanted rather than cells of human origin due to the immunosuppression required when using a xenograft method. This strategy allowed for comparison of host-transplant interactions and does not consider the differences in the developmental timelines between human and rodent nervous systems. Future studies transplanting human-derived tissues or organoids into various host rat ages would further elucidate the importance of host age on bi-directional connectivity between various combinations of host and transplant species and ages.

### Conclusion

Early cortical neural precursor cells grow, mature, and develop into a brain-like tissue after implantation at the CSF-SVZ brain interface rodents of up to one year of age at time of implantation. The newly formed BLT interacts with the host brain by extending long-distance neural processes into various host brain regions including the olfactory system, medial prefrontal cortex, and the striatum. The host brain structurally incorporates the BLT by supplying it with important support cells including astrocytes, oligodendrocytes, microglia, and blood vasculature.

The host also provides innervation input from the forebrain, thalamus, and striatum. Host age of up to one year does not affect growth and development of the BLT or any of the cellular components supplied by the host. Furthermore, the BLT projects into the host independent of age. As the age of the recipient at time of implantation increased, the long-distance host-to-graft innervation progressively decreased. With the rise of organoid brain transplantation to establish functional connectivity and modeling disease progression particularly at circuit level (Jgamadze et al., 2023; Revah et al., 2022), the influence of host age should become an important consideration. The ectopic placement of the transplant at the CSF tissue interface may enable easier assay of some host-transplant interactions. Finally, MRI was used to assess BLT growth, but it is likely that the use of MRI will grow in importance to assess functional integration and connectivity in cell and tissue transplantation studies non-invasively.

## Methods

### 1. Animal procedure

All animal procedures were handled according to the Institute of Laboratory Research guidelines and were approved by the Animal Care and Usen Committee (ACUC) of the National Institute of Neurological Disorders and Stroke.

### 2. Cell isolation and FACs sorting

E13.5 green fluorescent protein (GFP) embryos from Lewis rats were isolated according approved ACUC protocol and dorsal telencephalic region of developing cortical tissues were carefully dissected under magnifying scope. The tissues were dissociated using a papain-based enzyme. protocol, as previously described (Maric and Barker, Curr Protoc Neurosci 2005, PMID: 18428621). These cells were washed and maintained in neurobasal media prior to sorting. To purify the early neural stem/precursor cells, fluorescence activated cell sorting (FACS) was used to remove lineage-committed neuronal and glial progenitors and their post-mitotic counterparts, as well as to deplete non-neural cell phenotypes (microglia, endothelial cells, pericytes) from the (Lin^-^) neural stem/precursor population using a previously published method (Maric et al., Journal of Neuroscience 2007, PMID: 17314281). Briefly, telencephalic cell dissociates were stained using a panel of antibodies targeting the following surface markers: CD11b (to identify microglia), CD31 (to identify endothelial cells), NG2 proteoglycan (to identify pericytes), a cocktail of antibodies targeting A2B5, CD15 and CD24 (to identify neuroglial progenitor cells), GLAST (to identify radial glial cells and differentiating astrocytes), O4 (to identify oligodendroglia progenitors and their differentiating progeny), and a cocktail of antibodies targeting CD57, PSA-NCAM and GT1b gangliosides, via binding of tetanus toxin C-fragment (TnTx) and anti-TnTx antibody, to identify neuronal progenitors and differentiating neurons. Lin^-^ cortical NSCs were then purified by applying a lineage-dumping sorting protocol excluding all cell phenotypes expressing any of the markers listed above. Finally, a uniform single cell suspension of 5×10^4^ Lin^−^ cells/μL was prepared in neurobasal media (ThermoFisher Scientific, MA) supplemented with growth factors (bFGF and EGF, 20 ng/mL) and kept at 4 °C prior to the implantation.

### 3. Implantation into CSF-SVZ brain tissue interface near the rostral migratory stream (RMS)

21-to 24-days-old Lewis rats (40–60 g body weights) were used as recipients. The animals were anesthetized under 5% isoflurane in a 30% oxygen/70% nitrogen (oxygen-enhanced air) gas mixture. After anesthesia was induced, isoflurane level was adjusted to 2%. Under sterile conditions, a 1mm burr hole was drilled in the skulls of the animals (+1.5–1.6 AP and +1.5–1.6 ML from bregma and 3.8 DV from skull surface) above the lateral ventricles using stereotaxic coordinates from the Paxinos & Watson rat brain atlas. Cell suspension was loaded into a 10 μL glass syringe (Hamilton, MA) equipped with a 31-gauge needle. The needle was levered slowly and placed at a 4-mm depth from skull surface. Five microliter of cell suspension containing 5×10^4^ Lin^−^ cells/μL was slowly levered using a hand push over a 1-min period. After injection, the needle was left in situ for 3 min before removing. The burr hole was sealed with bone wax and the skin was sutured. Immediately after surgery, the animals were given analgesic (ketoprofen, 5 mg per kg). The animals were monitored for any sign of complication and returned to their home cages. For older hosts, 4-months and 12-months old rats were used. Coordinates were adjusted accordingly. For 4-month-old rats, coordinates were + 1.6-1.7 AP, +1.6-1.7 ML from Bregma and 4.2 DV from skull surface. For 12-months old rats, the coordinates were +1.7-1.8 AP, +1.7-1.8 ML and 4.4-4.5 DV from skull surface.

### 4. MRI

MRI was performed following implantation to monitor growth kinetics of the new tissue in the ventricle. All MRI experiments were done on an 11.7 T animal MRI system (30 cm 11.7 T horizontal magnet, Magnex Scientific, Oxford, England; MRI Electronics, Bruker Biospin, Billerica, MA) with a 12-cm integrated gradient shim system (Resonance Research Inc, Billerica, MA) using a custom-built volume transmit coil and a custom built, receive-only 2-coil array surface coil. Flash 3D gradient echo sequences were used for all MRI acquisitions. For in vivo imaging, the following parameters were used: field of view (FOV) = 1.92 cm3, matrix size 256 Å∼ 256 Å∼ 256 (100 μm isotropic resolution), 12.5 kHz bandwidth, TE = 8 ms, TR = 25 ms, and flip angle = 8°.

### 5. Immunostaining

For immunohistochemistry to characterize cell phenotypes, 10-μm thick brain coronal or sagittal sections were immunoreacted for 1 h at room temperature (RT) using 1 μg per mL final concentration (diluted in PBS supplemented with 1% bovine serum albumin, PBS/BSA) of the following primary antibodies (vendor source and product number are indicated in parentheses): mouse IgG1 anti-rat endothelial cell antigen (RECA1) (Abcam, ab9774), mouse IgG1 anti-CD68/ED1 (Thermo Fisher Scientific, MA5-16654), mouse IgG2a Olig2 (EMD Millipore, MABN50), mouse IgG2b anti-proliferation cell nuclear antigen (PCNA) (Abcam, ab184660), mouse IgG3 anti-glutamic acid decarboxylase 67 (GAD67) (Santa Cruz Biotechnology, sc-28376), mouse IgG2a anti-S100 (EMD Millipore, MAB079-1), anti-myelin basic protein (MBP) (EMD Millipore, MAB386), chicken IgY anti-glial fibrillary acidic protein (GFAP) (EMD Millipore, AB5541), rabbit IgG anti-Iba1 (Wako Chemicals, 019-19741), rabbit IgG anti-Doublecortin (DCX) (Abcam, ab18723), rabbit IgG anti-guinea pig IgG anti-NeuN (EMD Millipore, ABN90P). Select combinations of immunocompatible primary antibodies (i.e., antibodies from a different host or belonging to a different immunoglobulin class or subclass) from the list above were also used for multiplexing two or more biomarkers at the same time, as described previously (Maric et al., 2021) and further detailed in the results section. The sections were then washed in PBS/BSA and immunoreacted using a 1 μg per mL of the appropriate secondary antibodies (Thermo Fisher Scientific, Li-Cor Biosciences) conjugated to one of the following spectrally compatible fluorophores: Alexa Fluor 350, Alexa Fluor 405, Alexa Fluor 430, Alexa Fluor 488, Alexa Fluor 546, Alexa Fluor 594, Alexa Fluor 647, IRDye 680LT, or IRDye 800CW. Some sections were also counterstained with 1 μg per mL DAPI (Thermo Fisher Scientific) to facilitate cell counting. The slides with labeled tissue sections were then coverslipped using Immu-Mount medium (Thermo Fisher Scientific, MI). All sections were imaged using an Axio Imager Z.2 multi-channel scanning fluorescence microscope (Carl Zeiss, Thornwood, NY) equipped with a 20X Plan-Apochromat (Phase-2) objective Carl Zeiss), a high resolution ORCA-Flash 4.0 sCMOS digital camera (Hamamatsu Photonics, Japan) sensitive to a broad-spectrum of emission wavelengths, including those approaching infrared, a 200W X-Cite 200DC broad-spectrum light excitation source (Lumen Dynamics), and 10 self-contained excitation/dichroic/emission filter sets (Semrock, Rochester, NY) optimized to detect the following fluorophores with minimal spectral crosstalk: DAPI, Alexa Fluor 350, Alexa Fluor 405, Alexa Fluor 430, Alexa Fluor 488, Alexa Fluor 546, Alexa Fluor 594, Alexa Fluor 647, IRDye 680LT, and IRDye 800CW. Each labeling reaction was sequentially captured using filtered light through an appropriate fluorescence filter set and the images individually digitized at 16-bit resolution using the ZEN imaging program (Carl Zeiss). An appropriate color table was applied to each image to either match its emission spectrum or to set a distinguishing color balance. The pseudo-colored images were then converted into TIFF files, exported to Adobe Photoshop and overlaid as individual layers to create multi-colored merged composites.

For immunohistochemistry to assess development of long-range synapses, we used 30 μm thick brain coronal sections immunostained with anti-GFP and anti-OFP antibodies using a standard procedure for free-floating immunohistochemistry. Primary antibodies used for staining were mouse IgG1 monoclonal anti-mCherry (cross reacts with OFP) (Clonetech, 632543) and chicken polyclonal anti-GFP (Abcam, ab13970). These were visualized using Alexa Fluor 594 conjugated goat anti-mouse IgG1 and Alexa Fluor 488-conjugated goat anti-chicken IgY secondary antibodies (Thermo Fisher Scientific), respectively. Sections were mounted onto slides and coverslipped using Immu-Mount and images with a Nikon Eclipse Ti microscope (Nikon, CA) using a ×20 objective.

### 6. Immunogold labeling and electron microscopy

Rats were perfused according to the protocol described above but using 4% paraformaldehyde in PBS as fixative, and immunolabeled as described before (Tao-Cheng et al., 2021). Briefly, fixed brains were vibratomed into 100 μm thick coronal sections and processed free-floating in 24-well cell culture plates. Samples were made permeable and blocked with 0.1% saponin and 5% normal goat serum in PBS for 40–60 min, incubated with primary and then secondary antibodies (Nanogold, at 1:200, Nanoprobes, Yaphand, NY) for 1 h, fixed with 2% glutaraldehyde in PBS for 30 min, and stored at 4 °C in fixative. Samples were then silver enhanced (HQ kit, Nanoprobes), treated with 0.2% osmium tetroxide in 0.1 M phosphate buffer at pH 7.4 for 30 min on ice, then block stained with 0.25% uranyl acetate in acetate buffer at pH 5.0 for 1 h at 4 °C, dehydrated in graded ethanol, and embedded in epoxy resin.

### 7. Slice electrophysiology

To detect monosynaptic connections between the host and transplant, the ChR2-AAV used was: AAV2/9.hSynapsin.hChR2(H134R)-EYFP.WPRE.hGH, RRID: Addgene_26973 (Penn Vector Core, University of Pennsylvania). 8 weeks after transplant and 6 weeks after ChR2-AAV injection, rats were deeply anesthetized with 5% isoflurane vapors until the absence of righting reflex was observed. Animals were transcardially perfused with dissection buffer (80 mM NaCl, 3.5 mM KCl, 1.25 mM H2PO4, 25 mM NaHCO3, 4.5 mM MgSO4, 0.5 mM CaCl2, 10 mM glucose, and 90 mM sucrose) and the brain was removed. Tissue blocks were sectioned into 300 µm thick slices on Leica VT1000S vibratome (Leica Biosystems Inc.). Slices remained in the dissection buffer for 1-3 hours before being transferred to the submersion recording chamber continually perfused at 2 ml/min with artificial cerebral spinal fluid (ACSF; 124 mM NaCl, 5 mM KCl, 1.25 mM NaH2PO4·H2O, 26 mM NaHCO3, 10 mM dextrose, 2.5 mM CaCl2, and 1.5 mM MgCl2), which was bubbled with 95%/5% O2/CO2. The internal solution was K-gluconate based, which contained the following: 130 mM K-gluconate, 10 mM KCl, 0.2 mM EGTA, 10 mM HEPES, 4 mM MgATP, 0.5 mM NaGTP, and 10 mM Naphosphocreatine; pH 7.3, 280–290 mOsm). Cells with an access resistance higher than 25 MΩ and input resistance lower than 100 MΩ were discarded. An axon patch-clamp amplifier 700B (Molecular Devices) was used for voltage clamp recordings. Data were acquired through pClamp10 and analyzed with Clampfit 10.4 software (Molecular Devices). Neurons in the host or the transplant tissue were visually identified for recording. ChR2 was activated using a 455-nm light emitting diode (LED) DC2100, illuminated through the 40× objective lens and controlled by a digital stimulator (Cygnus DG4000A), both from ThorLabs. Cells which responded to LED stimulation with a short latency (<50 ms) were counted as responding monosynaptically. Neurons with no response, even to maximum LED intensity, were counted as not responding.

### 8. In vivo recording

The rats were anesthetized with urethane (1.25 g/kg, i.p. injection, Sigma-Aldrich). Once the absence of the hind-paw pinch reflexes was observed, the rat was mounted to a stereotaxic frame (Stoelting Inc.). An ocular ointment was applied to the eyes of the rat to prevent drying. Body temperature was kept constant at 37°C by using hand warmers and monitored via a thermometer throughout the entire recording sessions. Craniotomy (0.5-mm) was performed on the left orbitofrontal cortex. A 32-channel silicon probe (A1×32-6mm-50-177; NeuroNexus) was lowered to the target depth (#mm) from the brain surface. Ag/Ag-Cl pellet (EP1, World Precision Instruments) was inserted distal to the recording site as a ground and reference. Electrophysiological signals were sampled at 30 kHz (SmartBox Pro, NeuroNexus). Odor stimulation was delivered through a customized odor stimulator that used pneumatic pinch valves to provide the odor stimulus (10 % amyl acetate or mineral oil) without electrical noise. A vacuum system removed excess odorant effectively for the next trial and was balanced with the incoming air. One trial consisted of 800 ms odor stimulation and 1.5 s vacuum after 6 s from odor stimulus. Five trials were conducted with a 1 min inter-trial interval. Local field potential (LFP) data were analyzed using Spikes2 software (version 8.10a, Cambridge Electronic Design Limited). LFP signals were down-sampled at 1 kHz and band-pass filtered at 0.1-300 Hz. Odor stimulation evoked LFP signals were averaged and measured the LFP peak amplitude, onset time, and spectra.

### 9. AAV viral tracing

Eight to ten weeks post-implantation, AAV-ChR2-mCherry vectors were injected into the transplant tissue with MRI-guided coordinates. 300 μL of viral solution was injected at 50 μL/min using osmotic pump and waited 5 minutes after injection before the needle was retracted to minimize the backflow and leakage of the solution. The animals were euthanized for histological examination 2 weeks after the injection.

### 10. Image processing and quantification

To quantitatively assess the cell phenotype across the age groups, first, we manually delineated the regions of interest in both the BLT and host frontal cortex. Next, we utilized the Faster-RCNN method, as previously described, to locate cell nuclei using the DAPI and Histone channels. To classify the located cells into the major cell types of Neurons, Astrocytes, Microglia, and Oligodendrocytes, we utilized our cell classification model to extract a comprehensive representation of major cell types by employing the Capsule Network method (Maric et al., 2021). This abstract representation encapsulated information related to cell shape, morphology, and other features based on specific markers (NeuN, S100B, IBA1, Olig2). With the assistance of the Napari interactive viewer (Chiu et al., 2022). We performed thresholding on this abstract representation, to identify the major cell types in both the BLT and Cortex regions. We also measured the GFP intensity within bounding boxes and applied the thresholding technique to distinguish GFP^+^ cells. Further analysis within each cell class involved a comparison between GFP^+^ and GFP^-^ cells. Additionally, for blood vessels labeled with the RECA-1 biomarker, we employed the Segment Anything Model (Kirillov et al., 2023). This segmentation process was initiated by manually acquiring image samples and generating a segmentation mask. We calculated and compared the RECA-1 density between the BLT and host Cortex regions.

### 10. Statistical information

Data are expressed as mean ± standard error of the mean. Statistical analyses were performed using GraphPad (La Jolla, CA, USA). When evaluating cell percentages between the BLT and cortical regions, the comparison between the two groups was made using an unpaired T-test. Additionally, an unpaired T-test was used to compare the RECA1 percent area between the BLT and cortical region to evaluate vascular density. When comparing parameters between the three age groups, a one-way ANOVA and Tukey’s post-test for multiple comparisons was used. This was the analysis completed for the cell origin (GFP positive and GFP negative), anterograde projection density, and retrograde cell body density comparisons in which case all three age groups were directly compared.

## Data Availability

The datasets generated during and/or analyzed during the current study are available and will be deposited into Figshare.

## Acknowledgements

This work was supported by the Intramural Research Program of the National Institutes of Health (Grant number 1 Z1ANS003047-12), National Institute of Neurological Disorders and Stroke. The authors thank Dr. Patrick Wright for his assistance on preparation of figures and illustration, Francesca Venditti for assisting on analyzing part of the data, and Ms. Kathryn Sharer for obtaining rat embryos.

## Author contributions

N.P. designed and performed cell injection, MRI and fluorescent imaging experiments, wrote main manuscript text, and prepared figures.

G.G. designed and performed cell injection, viral tracing, MRI and fluorescent imaging, wrote main texts and prepare figures

D.M. performed precursor cell isolation and sorting and multiplex IHC staining and imaging experiments, wrote materials and method sections, and prepared the multiplex immunohistology figures.

J.-H.T.-C. performed EM experiments and prepared EM figures.

E.P. performed slice electrophysiology experiment and prepare figures

J. J. analyzed cell phenotype data and provided pipeline for analysis

H. J. performed in vivo recording

A.K. designed experiments.

All authors reviewed manuscript.

---

**Extended Figure 1.**
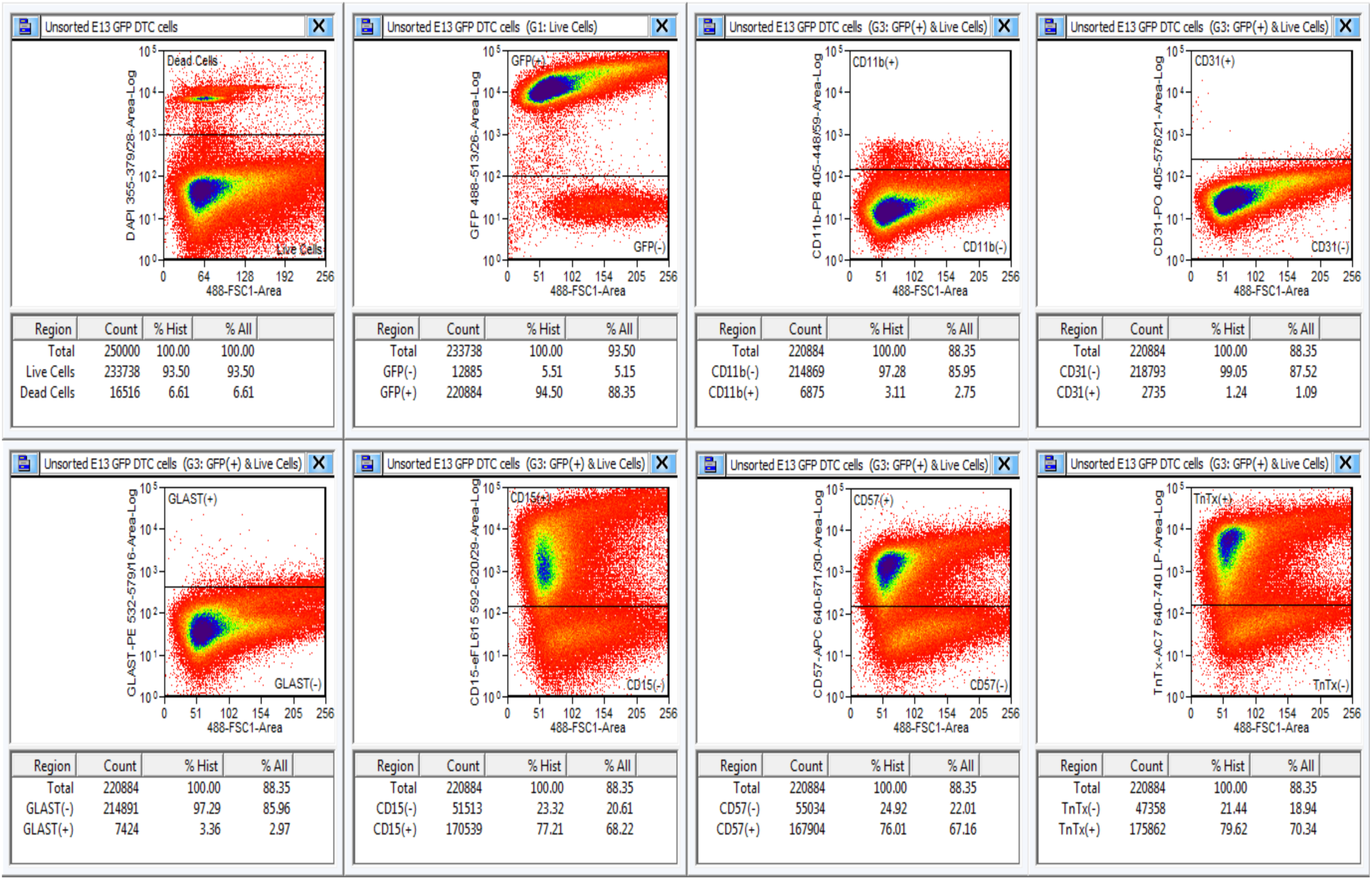
FACS isolation of GFP-expressing lineage-negative E13.5 cortical neural precursor cells using different surface markers to deplete microglia (CD11b), endothelial cells (CD31), radial glia and astroglial progenitors (GLAST), early neuroglial progenitors (CD15), and differentiating post-mitotic neurons (CD57, TnTx). Live and GFP-positive cells were selected using FACS. Of those, cells that were positive for surface markers were removed from the pool.

**Extended Figure 2.**
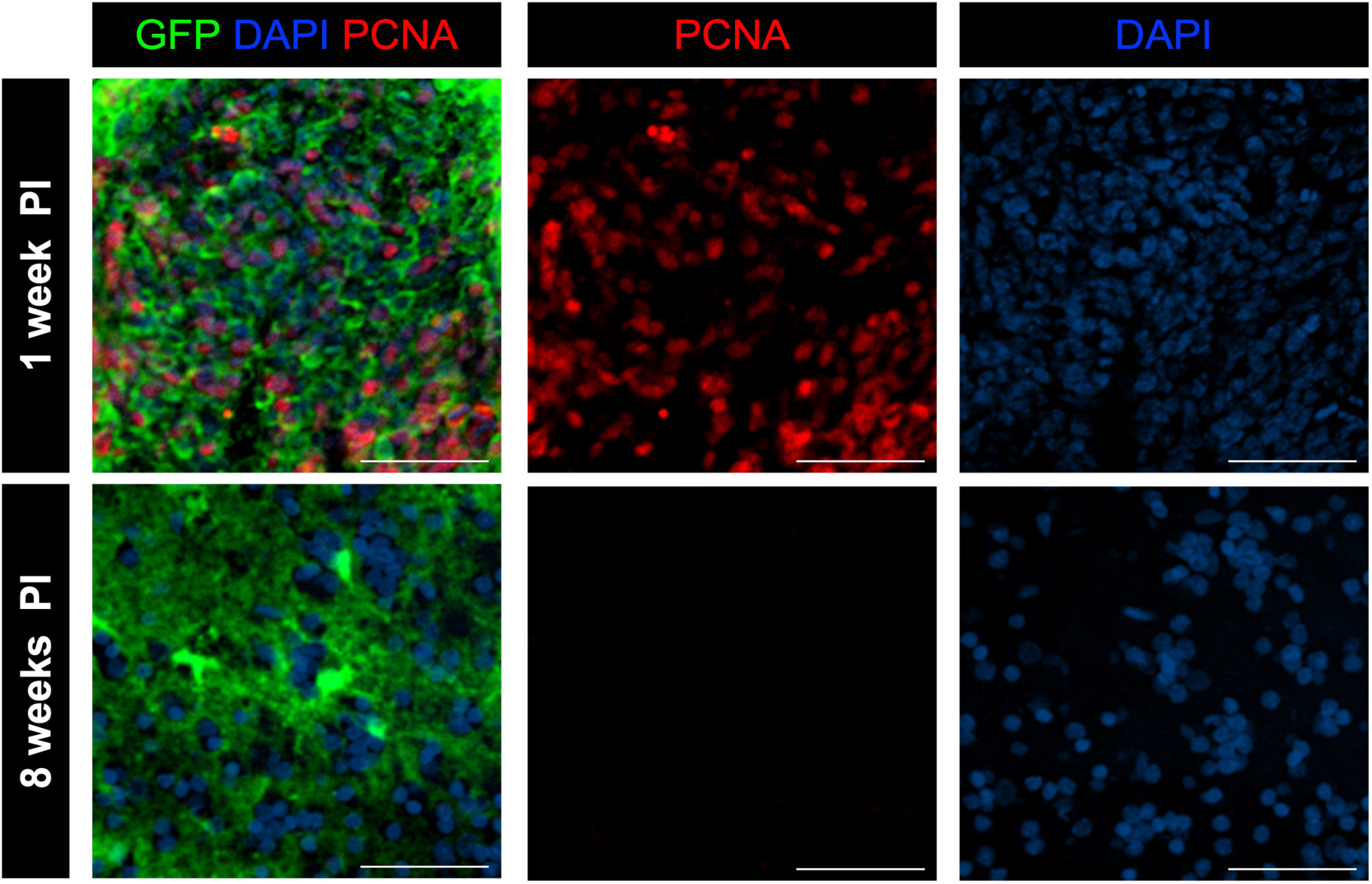
Immunostaining with PCNA shows that initially high proliferation activity within the precursor-derived BLT observed at 1-week post-implantation subsided by approximately 8 weeks post-implantation. Scale bar = 100 μm.

**Extended Figure 3.**
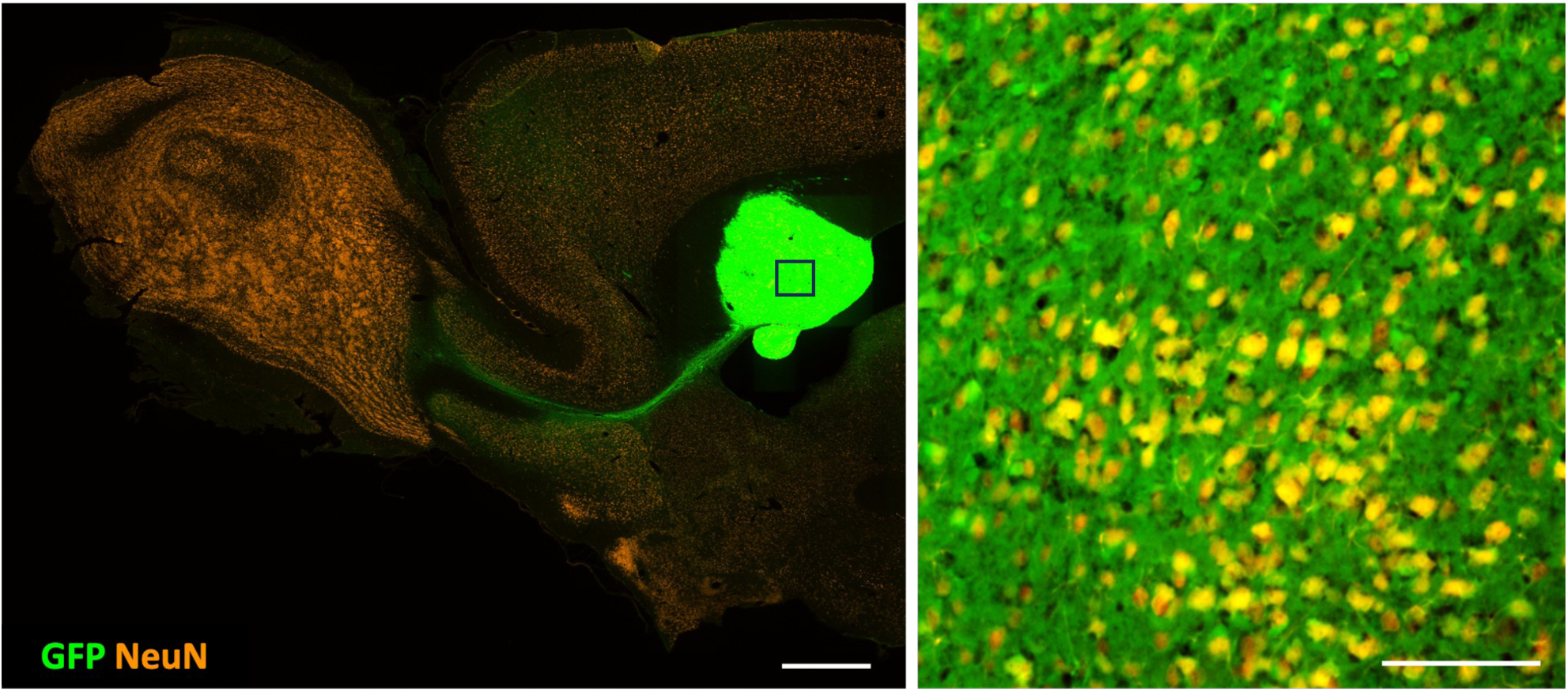
Sagittal fluorescent image of the host rat brain with the implanted BLT (green). BLT extended the outgrowth along the RMS pathway which is clear visible. Immunostaining with NeuN shows abundance of neurons that are mostly derived from the implanted GFP^+^ cells. Scale bar = 1 mm and 100 μm, respectively.

**Extended Figure 4.**
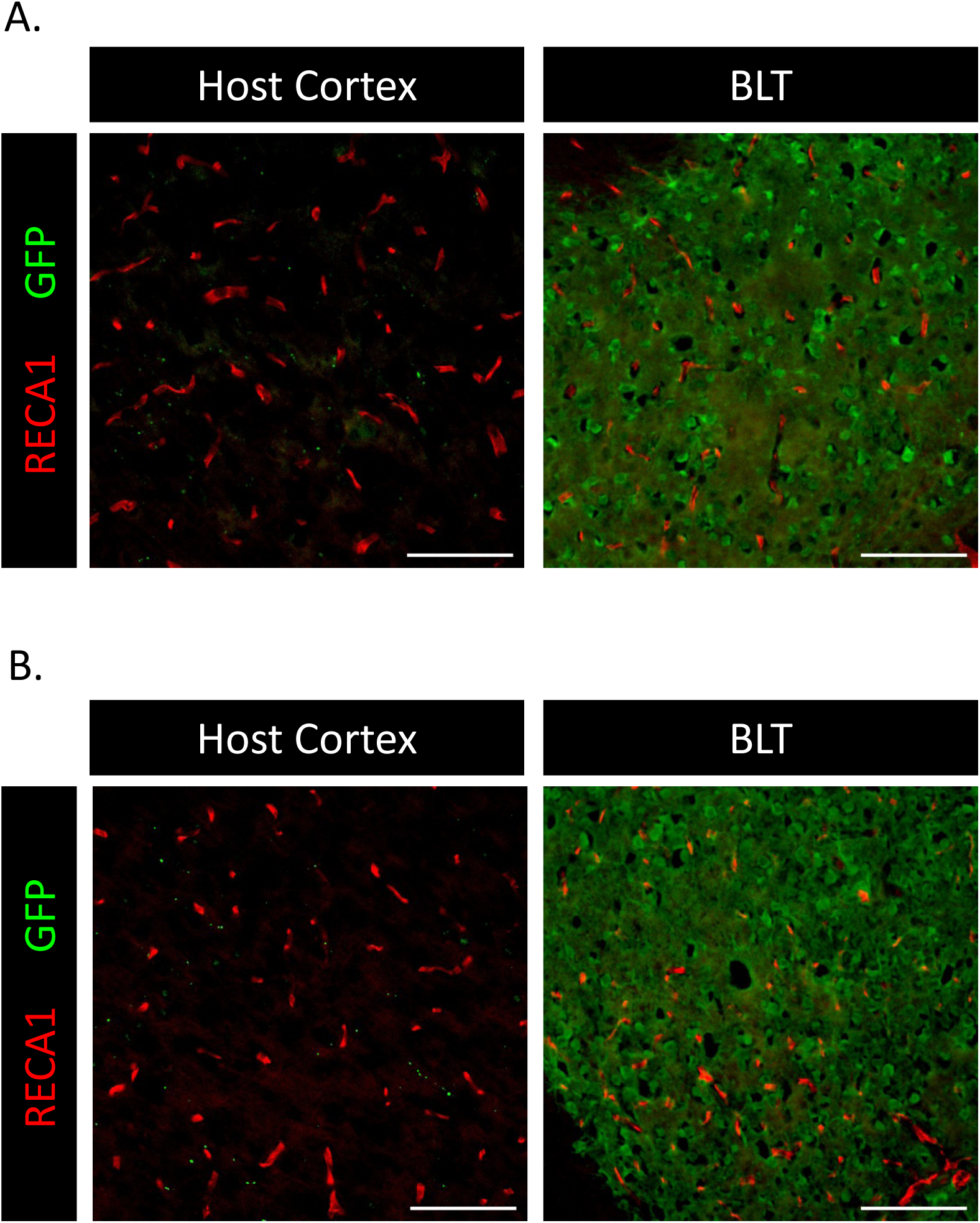
Blood vessels within the BLT and host cortex in 4-month-old (A) and 12-month-old host (B). RECA-1 immunostaining shows that the blood vessel density in the precursor-derived BLT remains similar across the host ages. Scale bar = 100 μm.

**Extended Figure 5.**
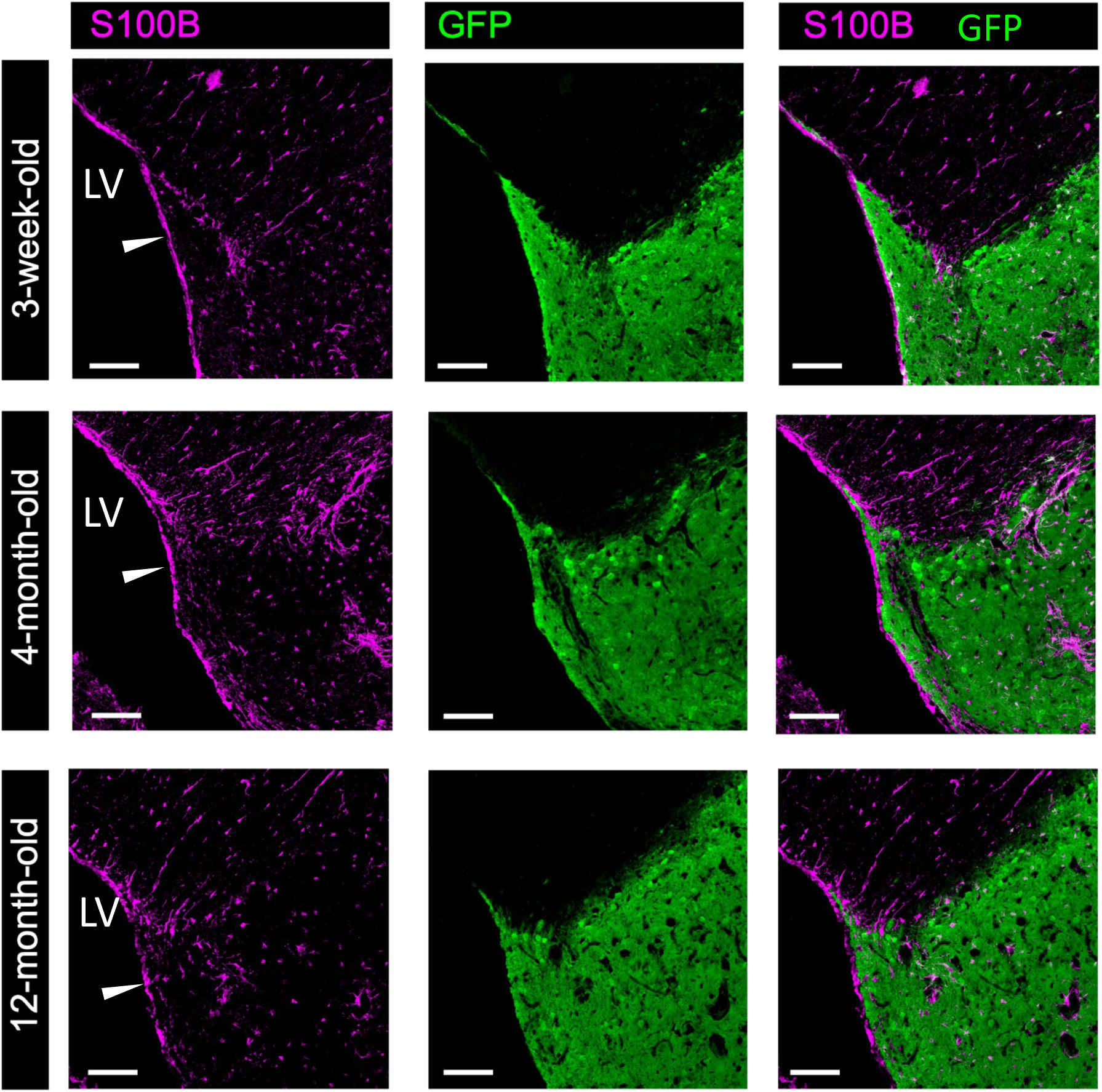
Immunostaining with S100B shows host ependymal cell layer (arrowhead) extending from the host brain parenchyma partially encapsulate the transplant GFP-positive tissue and separate this tissue from direct contact with the CSF-filled lateral ventricle (LV). Scale bar = 100 μm.

**Extended Figure 6.**
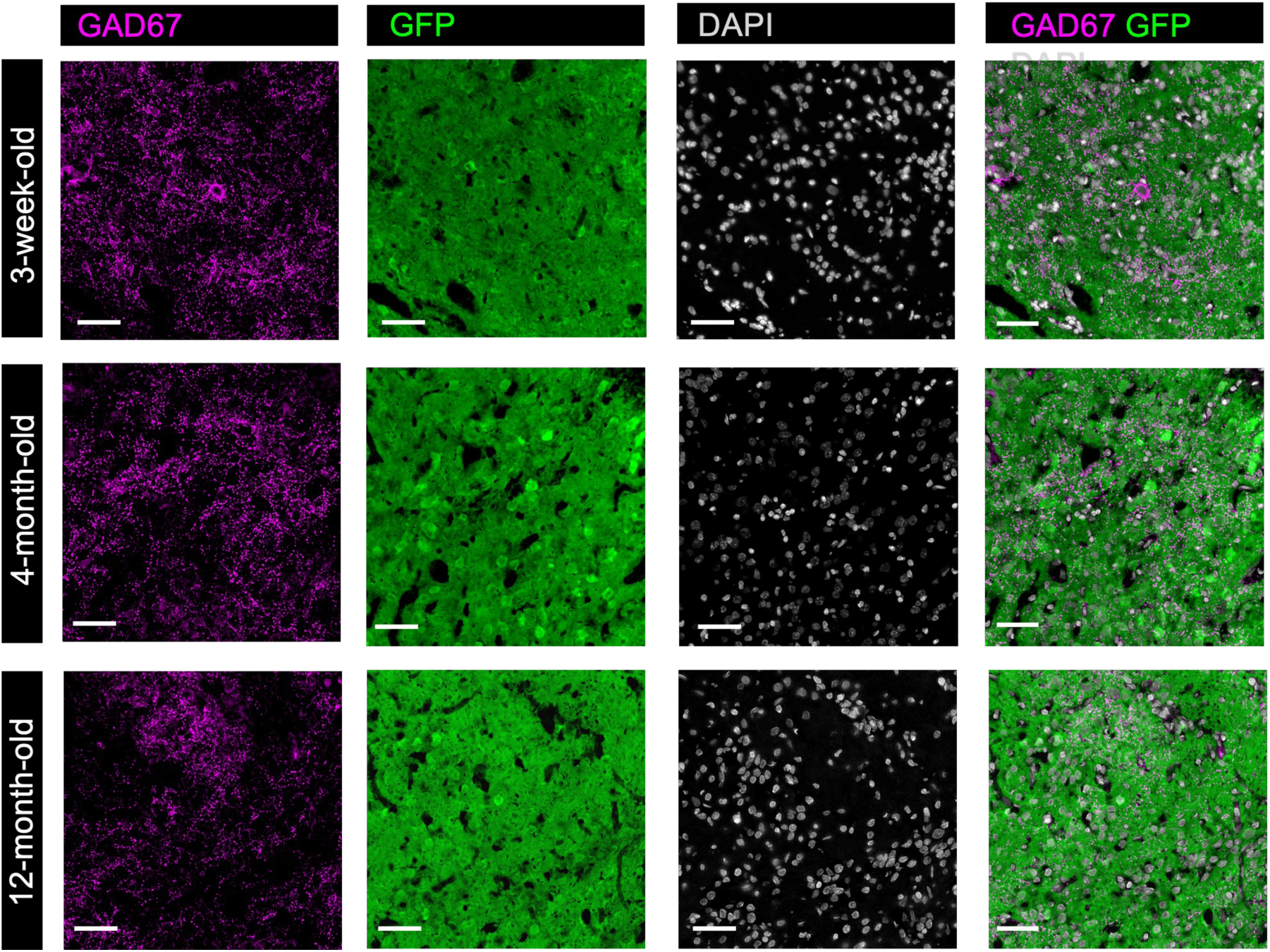
Presence of GAD67 (inhibitory input) positive processes and neurons within the BLT transplants. Immunostaining with GAD67 (magenta) show that these cells and processes are GFP-negative. Therefore, these cells have originated exclusively from the host. Scale bar = 50 μm.

**Extended Figure 7.**
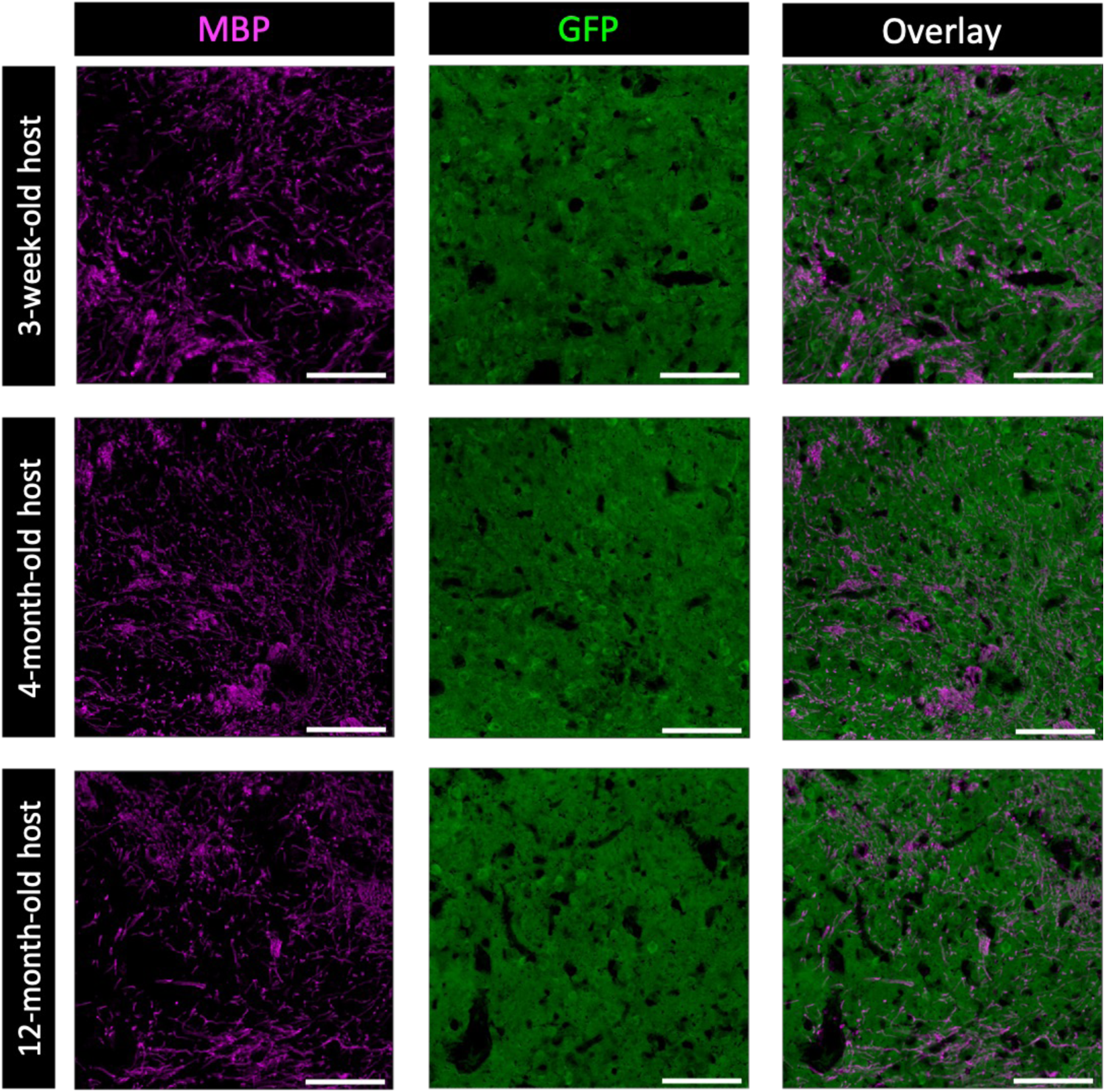
Immunostaining with myelin basic protein (MBP) indicates extensive myelination of BLT transplants across the host ages. MBP processes are also GFP-negative and thus have originated from the host. In good agreement with the previous report (Pothayee et al, 2018) based on electron microscopy and immunogold staining that the host myelinated the axons of the BLT-derived neurons. Scale bar = 100 μm.

**Extended Figure 8.**
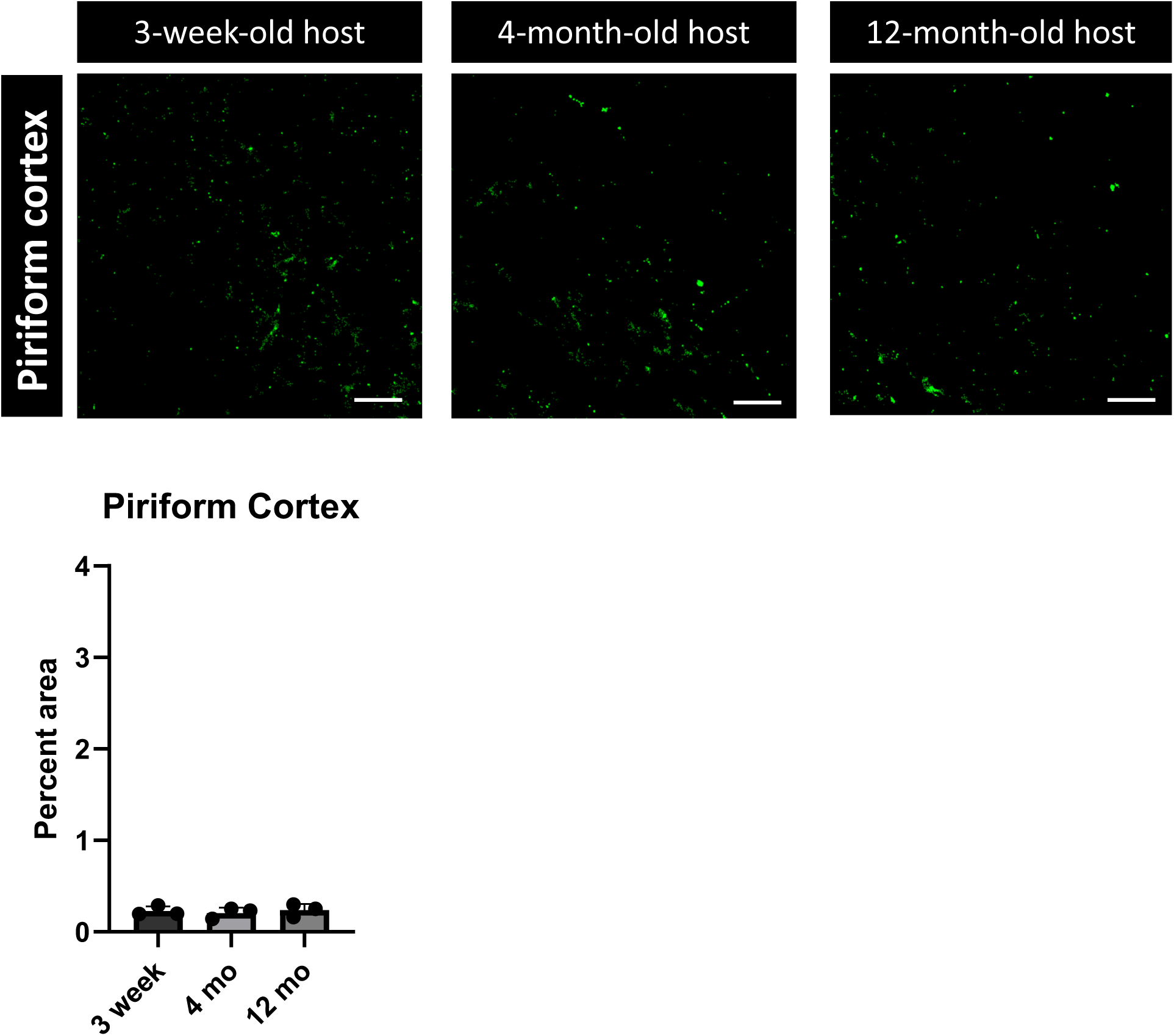
Fluorescent puncta and percentage of puncta per surface area depicting the extent of BLT projections in piriform cortex. Scale bar 100 μm.

## References

Akdemir, E. S., Huang, A. Y., & Deneen, B. (2020). Astrocytogenesis: where, when, and how. F1000Res, 9. 10.12688/f1000research.22405.1

Barker, R. A., Götz, M., & Parmar, M. (2018). New approaches for brain repair-from rescue to reprogramming. Nature, 557(7705), 329–334. 10.1038/s41586-018-0087-1

Chiu, C.-L., Clack, N., & community, t. n. (2022). napari: a Python Multi-Dimensional Image Viewer Platform for the Research Community. Microscopy and Microanalysis, 28(S1), 1576–1577. 10.1017/s1431927622006328

Denham, M., Parish, C. L., Leaw, B., Wright, J., Reid, C. A., Petrou, S., Dottori, M., & Thompson, L. H. (2012). Neurons derived from human embryonic stem cells extend long-distance axonal projections through growth along host white matter tracts after intra-cerebral transplantation. Front Cell Neurosci, 6, 11. 10.3389/fncel.2012.00011

Dulin, J. N., Adler, A. F., Kumamaru, H., Poplawski, G. H. D., Lee-Kubli, C., Strobl, H., Gibbs, D., Kadoya, K., Fawcett, J. W., Lu, P., & Tuszynski, M. H. (2018). Injured adult motor and sensory axons regenerate into appropriate organotypic domains of neural progenitor grafts. Nat Commun, 9(1), 84. 10.1038/s41467-017-02613-x

Espuny-Camacho, I., Michelsen, K. A., Gall, D., Linaro, D., Hasche, A., Bonnefont, J., Bali, C., Orduz, D., Bilheu, A., Herpoel, A., Lambert, N., Gaspard, N., Peron, S., Schiffmann, S. N., Giugliano, M., Gaillard, A., & Vanderhaeghen, P. (2013). Pyramidal neurons derived from human pluripotent stem cells integrate efficiently into mouse brain circuits in vivo. Neuron, 77(3), 440–456. 10.1016/j.neuron.2012.12.011

Espuny-Camacho, I., Michelsen, K. A., Linaro, D., Bilheu, A., Acosta-Verdugo, S., Herpoel, A., Giugliano, M., Gaillard, A., & Vanderhaeghen, P. (2018). Human Pluripotent Stem-Cell-Derived Cortical Neurons Integrate Functionally into the Lesioned Adult Murine Visual Cortex in an Area-Specific Way. Cell Rep, 23(9), 2732–2743. 10.1016/j.celrep.2018.04.094

Falkner, S., Grade, S., Dimou, L., Conzelmann, K. K., Bonhoeffer, T., Gotz, M., & Hubener, M. (2016). Transplanted embryonic neurons integrate into adult neocortical circuits. Nature, 539(7628), 248–253. 10.1038/nature20113

Feigin, V. L., Vos, T., Nichols, E., Owolabi, M. O., Carroll, W. M., Dichgans, M., Deuschl, G., Parmar, P., Brainin, M., & Murray, C. (2020). The global burden of neurological disorders: translating evidence into policy. Lancet Neurol, 19(3), 255–265. 10.1016/s1474-4422(19)30411-9

Fuchs, E., Tumbar, T., & Guasch, G. (2004). Socializing with the neighbors: stem cells and their niche. Cell, 116(6), 769–778. 10.1016/s0092-8674(04)00255-7

Gaillard, A., Prestoz, L., Dumartin, B., Cantereau, A., Morel, F., Roger, M., & Jaber, M. (2007). Reestablishment of damaged adult motor pathways by grafted embryonic cortical neurons. Nat Neurosci, 10(10), 1294–1299. 10.1038/nn1970

Gash, D. M., Collier, T. J., & Sladek, J. R., Jr. (1985). Neural transplantation: a review of recent developments and potential applications to the aged brain. Neurobiol Aging, 6(2), 131–174. 10.1016/0197-4580(85)90031-4

Girman, S. V., & Golovina, I. L. (1990). Electrophysiological properties of embryonic neocortex transplants replacing the primary visual cortex of adult rats. Brain Res, 523(1), 78–86. 10.1016/0006-8993(90)91637-v

Grade, S., & Götz, M. (2017). Neuronal replacement therapy: previous achievements and challenges ahead. NPJ Regen Med, 2, 29. 10.1038/s41536-017-0033-0

Gronning Hansen, M., Laterza, C., Palma-Tortosa, S., Kvist, G., Monni, E., Tsupykov, O., Tornero, D., Uoshima, N., Soriano, J., Bengzon, J., Martino, G., Skibo, G., Lindvall, O., & Kokaia, Z. (2020). Grafted human pluripotent stem cell-derived cortical neurons integrate into adult human cortical neural circuitry. Stem Cells Transl Med, 9(11), 1365–1377. 10.1002/sctm.20-0134

Harel, N. Y., & Strittmatter, S. M. (2006). Can regenerating axons recapitulate developmental guidance during recovery from spinal cord injury? Nat Rev Neurosci, 7(8), 603–616. 10.1038/nrn1957

Jgamadze, D., Lim, J. T., Zhang, Z., Harary, P. M., Germi, J., Mensah-Brown, K., Adam, C. D., Mirzakhalili, E., Singh, S., Gu, J. B., Blue, R., Dedhia, M., Fu, M., Jacob, F., Qian, X., Gagnon, K., Sergison, M., Fruchet, O., Rahaman, I.,…Chen, H. I. (2023). Structural and functional integration of human forebrain organoids with the injured adult rat visual system. Cell Stem Cell, 30(2), 137–152 e137. 10.1016/j.stem.2023.01.004

Kelley, K. W., Revah, O., Gore, F., Kaganovsky, K., Chen, X., Deisseroth, K., & Pașca, S. P. (2024). Host circuit engagement of human cortical organoids transplanted in rodents. Nat Protoc. 10.1038/s41596-024-01029-4

Kirillov, A., Mintun, E., Ravi, N., Mao, H., Rolland, C., Gustafson, L., Xiao, T., Whitehead, S., Berg, A. C., Lo, W.-Y., Dollár, P., & Girshick, R. B. (2023). Segment Anything. 2023 IEEE/CVF International Conference on Computer Vision (ICCV), 3992-4003.

Kitahara, T., Sakaguchi, H., Morizane, A., Kikuchi, T., Miyamoto, S., & Takahashi, J. (2020). Axonal Extensions along Corticospinal Tracts from Transplanted Human Cerebral Organoids. Stem Cell Reports, 15(2), 467–481. 10.1016/j.stemcr.2020.06.016

Kumamaru, H., Lu, P., Rosenzweig, E. S., Kadoya, K., & Tuszynski, M. H. (2019). Regenerating Corticospinal Axons Innervate Phenotypically Appropriate Neurons within Neural Stem Cell Grafts. Cell Rep, 26(9), 2329–2339 e2324. 10.1016/j.celrep.2019.01.099

Lu, P., Wang, Y., Graham, L., McHale, K., Gao, M., Wu, D., Brock, J., Blesch, A., Rosenzweig, E. S., Havton, L. A., Zheng, B., Conner, J. M., Marsala, M., & Tuszynski, M. H. (2012). Long-distance growth and connectivity of neural stem cells after severe spinal cord injury. Cell, 150(6), 1264–1273. 10.1016/j.cell.2012.08.020

Lu, S. Y., & Norman, A. B. (1993). Neurotransmitter receptors in fetal tissue transplants: expression and functional significance. J Neural Transplant Plast, 4(3), 215–226. 10.1155/np.1993.215

Mansour, A. A., Goncalves, J. T., Bloyd, C. W., Li, H., Fernandes, S., Quang, D., Johnston, S., Parylak, S. L., Jin, X., & Gage, F. H. (2018). An in vivo model of functional and vascularized human brain organoids. Nat Biotechnol, 36(5), 432–441. 10.1038/nbt.4127

Maric, D., Fiorio Pla, A., Chang, Y. H., & Barker, J. L. (2007). Self-renewing and differentiating properties of cortical neural stem cells are selectively regulated by basic fibroblast growth factor (FGF) signaling via specific FGF receptors. J Neurosci, 27(8), 1836–1852. 10.1523/JNEUROSCI.5141-06.2007

Maric, D., Jahanipour, J., Li, X. R., Singh, A., Mobiny, A., Van Nguyen, H., Sedlock, A., Grama, K., & Roysam, B. (2021). Whole-brain tissue mapping toolkit using large-scale highly multiplexed immunofluorescence imaging and deep neural networks. Nat Commun, 12(1), 1550. 10.1038/s41467-021-21735-x

Michelsen, K. A., Acosta-Verdugo, S., Benoit-Marand, M., Espuny-Camacho, I., Gaspard, N., Saha, B., Gaillard, A., & Vanderhaeghen, P. (2015). Area-specific reestablishment of damaged circuits in the adult cerebral cortex by cortical neurons derived from mouse embryonic stem cells. Neuron, 85(5), 982–997. 10.1016/j.neuron.2015.02.001

Palma-Tortosa, S., Tornero, D., Gronning Hansen, M., Monni, E., Hajy, M., Kartsivadze, S., Aktay, S., Tsupykov, O., Parmar, M., Deisseroth, K., Skibo, G., Lindvall, O., & Kokaia, Z. (2020). Activity in grafted human iPS cell-derived cortical neurons integrated in stroke-injured rat brain regulates motor behavior. Proc Natl Acad Sci U S A, 117(16), 9094–9100. 10.1073/pnas.2000690117

Pothayee, N., Maric, D., Sharer, K., Tao-Cheng, J. H., Calac, A., Bouraoud, N., Pickel, J., Dodd, S., & Koretsky, A. (2018). Neural precursor cells form integrated brain-like tissue when implanted into rat cerebrospinal fluid. Commun Biol, 1, 114. 10.1038/s42003-018-0113-8

Price, D. J., Kennedy, H., Dehay, C., Zhou, L., Mercier, M., Jossin, Y., Goffinet, A. M., Tissir, F., Blakey, D., & Molnár, Z. (2006). The development of cortical connections. Eur J Neurosci, 23(4), 910–920. 10.1111/j.1460-9568.2006.04620.x

Quezada, A., Ward, C., Bader, E. R., Zolotavin, P., Altun, E., Hong, S., Killian, N. J., Xie, C., Batista-Brito, R., & Hebert, J. M. (2023). An In Vivo Platform for Rebuilding Functional Neocortical Tissue. Bioengineering (Basel), 10(2). 10.3390/bioengineering10020263

Revah, O., Gore, F., Kelley, K. W., Andersen, J., Sakai, N., Chen, X., Li, M. Y., Birey, F., Yang, X., Saw, N. L., Baker, S. W., Amin, N. D., Kulkarni, S., Mudipalli, R., Cui, B., Nishino, S., Grant, G. A., Knowles, J. K., Shamloo, M.,…Pasca, S. P. (2022). Maturation and circuit integration of transplanted human cortical organoids. Nature, 610(7931), 319–326. 10.1038/s41586-022-05277-w

Varadarajan, S. G., Hunyara, J. L., Hamilton, N. R., Kolodkin, A. L., & Huberman, A. D. (2022). Central nervous system regeneration. Cell, 185(1), 77–94. 10.1016/j.cell.2021.10.029

Velasco, S., Kedaigle, A. J., Simmons, S. K., Nash, A., Rocha, M., Quadrato, G., Paulsen, B., Nguyen, L., Adiconis, X., Regev, A., Levin, J. Z., & Arlotta, P. (2019). Individual brain organoids reproducibly form cell diversity of the human cerebral cortex. Nature, 570(7762), 523–527. 10.1038/s41586-019-1289-x

Wang, M., Zhang, L., Novak, S. W., Yu, J., Gallina, I. S., Xu, L. L., Lim, C. K., Fernandes, S., Shokhirev, M. N., Williams, A. E., Saxena, M. D., Coorapati, S., Parylak, S. L., Quintero, C., Molina, E., Andrade, L. R., Manor, U., & Gage, F. H. (2024). Morphological diversification and functional maturation of human astrocytes in glia-enriched cortical organoid transplanted in mouse brain. Nat Biotechnol. 10.1038/s41587-024-02157-8

Babcock, K. R., Page, J. S., Fallon, J. R., & Webb, A. E. (2021). Adult Hippocampal Neurogenesis in Aging and Alzheimer’s Disease. Stem Cell Reports, 16(4), 681–693. 10.1016/j.stemcr.2021.01.019

Couillard-Despres, S., Iglseder, B., & Aigner, L. (2011). Neurogenesis, cellular plasticity and cognition: the impact of stem cells in the adult and aging brain--a mini-review. Gerontology, 57(6), 559–564. 10.1159/000323481

Molnár, Z., & Blakemore, C. (1995). How do thalamic axons find their way to the cortex? Trends Neurosci, 18(9), 389–397. 10.1016/0166-2236(95)93935-q

Su, Y., Wang, X., Yang, Y., Chen, L., Xia, W., Hoi, K. K., Li, H., Wang, Q., Yu, G., Chen, X., Wang, S., Wang, Y., Xiao, L., Verkhratsky, A., Fancy, S. P. J., Yi, C., & Niu, J. (2023). Astrocyte endfoot formation controls the termination of oligodendrocyte precursor cell perivascular migration during development. Neuron, 111(2), 190–201.e198. 10.1016/j.neuron.2022.10.032

Talaverón, R., Matarredona, E. R., de la Cruz, R. R., Macías, D., Gálvez, V., & Pastor, A. M. (2014). Implanted neural progenitor cells regulate glial reaction to brain injury and establish gap junctions with host glial cells. Glia, 62(4), 623–638. 10.1002/glia.22630

Tao-Cheng, J. H., Crocker, V., Moreira, S. L., & Azzam, R. (2021). Optimization of protocols for pre-embedding immunogold electron microscopy of neurons in cell cultures and brains. Mol Brain, 14(1), 86. 10.1186/s13041-021-00799-2

